# A high-quality chromosome-level genome assembly of rohu carp, *Labeo rohita*, and its utilization in SNP-based exploration of gene flow and sex determination

**DOI:** 10.1101/2022.09.08.507226

**Authors:** Mark A. Arick, Corrinne E. Grover, Chuan-Yu Hsu, Zenaida Magbanua, Olga Pechanova, Emma R. Miller, Adam Thrash, Ramey C. Youngblood, Lauren Ezzell, Md Samsul Alam, John A. H. Benzie, Matthew G. Hamilton, Attila Karsi, Mark L. Lawrence, Daniel G. Peterson

## Abstract

*Labeo rohita* (rohu) is a carp important to aquaculture in South Asia, with a production volume close to Atlantic salmon. While genetic improvements to rohu are ongoing, the genomic methods commonly used in other aquaculture improvement programs have historically been precluded in rohu, partially due to the lack of a high quality reference genome. Here we present a high-quality *de novo* genome produced using a combination of next-generation sequencing technologies, resulting in a 946 Mb genome consisting of 25 chromosomes and 2,844 unplaced scaffolds. Notably, while approximately half the size of the existing genome sequence, our genome represents 97.9% of the genome size newly estimated here using flow cytometry. Sequencing from 120 individuals was used in conjunction with this genome to predict the population structure, diversity, and divergence in three major rivers (Jamuna, Padma, and Halda), in addition to infer a likely sex determination mechanism in rohu. These results demonstrate the utility of the new rohu genome in modernizing some aspects of rohu genetic improvement programs.

## Introduction

*Labeo rohita* (rohu; rui), a carp naturally found in the Indo-Gangetic and surrounding river systems (Das et al. 2020), is an important aquaculture fish in many areas of South Asia (FAO 2020). The annual aquaculture production of rohu in Bangladesh was 386.3 thousand tonnes in the 2019-2020 fiscal year, the second-highest among all aquaculture species in the country (DoF 2020). Annual aquaculture production of the species is approximately 2.0 million metric tons (Mt) globally, a volume comparable with *Salmo salar* (Atlantic salmon; 2.4 Mt); however, study and understanding of rohu genomics is not commensurate with its global significance (Rasal and Sundaray 2020). Although there is increasing interest in applying next-generation sequencing (NGS) and other high-throughput methods to rohu (Robinson et al. 2014; Rasal et al. 2017; Hamilton et al. 2019; Rasal et al. 2020; Sahoo et al. 2021), to date, most studies have been conducted in the absence of a genome sequence. Recently, a draft genome was published for rohu (Das et al. 2020) to provide a unifying resource for NGS analysis; however, the relatively low quality of the genome limits the development of a robust genomic framework for the species.

Genetically improved rohu seed is increasingly available to farmers, from both mass-selection (e.g. ‘Subarna Rohu’ in Bangladesh and ‘Ayeyarwady Hatchery’ in Myanmar) (Hamilton 2019; SZA 2021 Jun 10) and family-based (i.e. pedigree-based) improvement programs (e.g. ‘Jayanti’ in India and ‘WorldFish Genetically Improved Rohu’ in Bangladesh) (Das Mahapatra et al. 2007; Rasal et al. 2017; Hamilton et al. 2019; Hamilton et al. 2022). However, genomic methods routinely applied in other aquaculture species (e.g., parentage assignment and genomic selection) have yet to be routinely applied in rohu genetic improvement programs (Sahoo et al. 2017; Rasal and Sundaray 2020), primarily due to a historical focus on improving growth rate (directly assessable at low cost on selection candidates), limited financial resources, and the absence of a genome sequence. As existing family-based programs expand to include additional traits (e.g., carcass traits, feed conversion ratio, tolerance to extreme environments, and disease resistance) (Rasal and Sundaray 2020), the advantages afforded by improved genomic resources in rohu will become increasingly compelling.

The mechanism of sex determination (SD) in rohu is a lingering question with applications to aquaculture, as understanding SD mechanisms in other species has been used to prevent precocious maturation, exploit sexual dimorphism in growth rate, improve carcass quality, and protect both environmental values and intellectual property (Budd et al. 2015). Despite its relevance to aquaculture and genetic improvement, SD in rohu has been understudied (Sahoo et al. 2021) both due to the high diversity of teleost SD mechanisms (Heule et al. 2014) and the lack of high-quality genomic resources (Sahu et al. 2013).

Here we present a new *de novo* high-quality genome for rohu that improves sequence contiguity and reduces duplication. We use this reference to assess diversity among populations of rohu from three different rivers and to preliminarily describe the gametic system of sex determination in rohu, demonstrating the utility of this improved sequence to increase understanding and facilitate aquacultural production and genetic improvement.

## Methods & Materials

### Sample Collection, DNA Extraction, & Sequencing

Blood samples were collected from five individual rohu (hence referred to as Rohu-1 through Rohu-5) from a fish farm located in the District of Rangpur, Bangladesh. The fish were handled as per guidelines of the Ethics Standard Review Committee of Bangladesh Agricultural University (BAU) involving fish and animals (Approval No. BAURES/ESRC/2019/Fish/01). Each fish was euthanized using clove oil, dissected, and blood was collected from the heart using a syringe. Each blood sample was placed in an ethylenediaminetetraacetic acid (EDTA) containing vial, and vials were shipped in an insulated container to Mississippi State University (MSU) for DNA extraction.

High molecular weight (HMW) genomic DNA for whole genome sequencing was extracted from 150 μL of blood from Rohu-1 using CTAB lysis buffer followed by the phenol/chloroform purification procedure (Doyle and Doyle 1987). The concentration and purity of extracted genomic DNA samples were measured by a NanoDrop ND-1000 spectrophotometer (NanoDrop Technologies, Wilmington, DE, USA). The quality of genomic DNA was validated by electrophoresis on a 0.8% w/v agarose gel.

The genomic DNA from Rohu-1 was used to prepare 10 Oxford Nanopore R9.4 MinION flow cells. For each flow cell, 2 to 2.5 μg of genomic DNA and a Nanopore Genomic DNA Ligation Sequencing Kit SQK-LSK 109 (Oxford Nanopore Technologies, Oxford, UK) were used to create a DNA library. For each of the 10 libraries, 700 to 750 ng of DNA was loaded onto a Nanopore Flow Cell R9.4.1 (Oxford Nanopore Technologies, Oxford, UK) and sequenced on a GridION sequencer (Oxford Nanopore Technologies, Oxford, UK) for 48 hours.

Rohu-1 genomic DNA was also sequenced on an Illumina HiSeq X-Ten (2×150 bp). In brief, 2 μg of Rohu-1 genomic DNA was used with an Illumina TruSeq DNA PCR-free Library Prep Kit (Illumina, San Diego, CA, USA) to create an Illumina sequencing library. The final DNA-Seq library, which had an insert size range of 350 bp to 450 bp, was submitted to Novogene (www.en.novogene.com) for two lanes of PE150 on an Illumina HiSeq X-Ten (Illumina, San Diego, CA, USA) sequencer.

A Hi-C library also was prepared using 100 μL of Rohu-1 blood with the Proximo Hi-C Animal Kit (Phase Genomics, Seattle, WA, USA). The final Hi-C DNA-Seq library was submitted to Novogene (www.en.novogene.com) for one lane of PE150 Illumina HiSeq X-Ten (Illumina, San Diego, CA, USA) sequencing.

Lastly, Rohu-1 blood cells were embedded in agarose and high molecular weight DNA was isolated according to the Bionano Prep Frozen Blood Protocol (Bionano Genomics, San Diego, CA). The extracted DNA molecules were labeled with the Direct Label and Stain (DLS) DNA Labeling kit (Bionano Genomics, San Diego, CA). Once labeled and stained, the DNA was imaged on the Bionano Saphyr instrument (Bionano Genomics, San Diego, CA).

### Genome Size Estimation

The genome size of rohu was estimated using two independent methods: flow cytometry and k-mer profiling.

Flow cytometry was performed using erythrocyte nuclei from Rohu-1, Rohu-2, Rohu-3, Rohu-4, and Rohu-5 using trout erythrocyte nuclei (TENs; https://www.biosure.com/tens.html) as a standard (1C = 6.5 pg). For each replicate, nuclei were stabilized in 200 μl of LB01-propidium iodide (PI) buffer as per (Pellicer and Leitch 2014), and two drops of TENs standard were used per 50μl of fish blood. Each sample was measured twice, totaling 10 runs overall. Only measurements with greater than 5,000 nuclei and a coefficient of variation (CV) of less than 3% were retained (Pellicer and Leitch 2014).

For k-mer profiling, Jellyfish [v2.2.10] (Marçais and Kingsford 2011) was used to ‘‘digest” the Rohu-1 Illumina paired reads into 50-mers. GenomeScope [v1.0] (Vurture et al. 2017) was then used to estimate genome size using the resulting k-mer profile.

### Assembly & Annotation

Nanopore sequence data was filtered to remove the control lambda-phage and sequences shorter than 1000 bases using the nanopack tool suite [v1.0.1] (De Coster et al. 2018). Trimmomatic [v0.32] (Bolger et al. 2014) was used to remove adapters, trim low-quality bases, and filter out reads shorter than 85bp. The filtered nanopore data were assembled into contigs using wtdbg2 [v2.4] (Ruan and Li 2020). The contigs were polished using two iterations of racon [v1.4.0] (Vaser et al. 2017) with minimap2 [v2.17] (Li 2018) mapping the nanopore reads. The contigs were further polished with Illumina paired end read data using pilon [v1.23] (Walker et al. 2014) with bwa [v0.7.10] (Li 2013 May 26) mapping the Illumina paired reads. The resulting contigs were scaffolded using Bionano Solve [Solve3.4.1_09262019] and SALSA [v2.3] (Ghurye et al. 2019). Those scaffolds larger than 10Mb were linked and oriented based on the *Onychostoma macrolepis* genome (Sun et al. 2020), the chromosome assembly most similar to rohu available on NCBI, using RagTag [v1.1.1] (Alonge et al. 2021).

RepeatModeler [v2.0.1] (Flynn et al. 2020) and RepeatMasker [v4.1.1] (Smit et al. 2013 2015) were used to create a species-specific repeat database, and this database was subsequently used by RepeatMasker to mask those repeats in the genome. All available RNA-seq libraries for rohu (comprising brain, pituitary, gonad, liver, pooled, and whole body tissues for both sexes; Supplemental Table 1) were downloaded from NCBI and mapped to the masked genome using hisat2 [v2.1.0] (Kim et al. 2019). These alignments were used in both the mikado [v2.0rc2] (Venturini et al. 2018) and braker2 [v2.1.5] (Brůna et al. 2021) pipelines. Mikado uses putative transcripts assembled from the RNA-seq alignments generated via stringtie [v2.1.2] (Kovaka et al. 2019), cufflinks [v2.2.1] (Trapnell et al. 2012), and trinity [v2.11.0] (Grabherr et al. 2011) along with the junction site prediction from portcullis [v1.2.2] (Mapleson et al. 2018), the alignments of the putative transcripts with UniprotKB Swiss-Prot [v2021.03] (The UniProt Consortium 2021), and the ORFs from prodigal [v2.6.3] (Hyatt et al. 2010) to select the best representative transcript for each locus. Braker2 uses those RNA-seq alignments and the gene prediction from GeneMark-ES [v4.61] (Borodovsky and Lomsadze 2011) to train a species-specific Augustus [v3.3.3] (Stanke et al. 2006) model. Maker2 [v2.31.10] (Holt and Yandell 2011) predicts genes based on the new Augustus, GeneMark, and SNAP models derived from Braker2, modifying the predictions based on the available RNA and protein evidence from the Cyprinidae family in the NCBI RefSeq database. Any predicted genes with an annotation edit distance (AED) above 0.47 were removed from further analysis. The remaining genes were functionally annotated using InterProScan [v5.47-82.0] (Jones et al. 2014) and BLAST+ [v2.9.0] (Camacho et al. 2009) alignments against the UniprotKB Swiss-Prot database. BUSCO [v5.2.2] (Manni et al. 2021) was used to verify the completeness of both the genome and annotations against the actinopterygii_odb10 database. Lastly, genes spanning large gaps or completely contained within another gene on the opposite strand were removed using a custom Perl script (https://github.com/IGBB/rohu-genome/).

### Comparative Genomics

The assembly statistics, length distributions, BUSCO completeness scores, and sequence similarity via dot-plots were compared between the IGBB rohu genome (reported here) and the rohu genome reported by (Das et al. 2020) (CIFA, Refseq accession GCA_004120215.1), as well as all 12 annotated Cypriniformes genomes from NCBI (Table 1). Assembly statistics were calculated using abyss-fac from ABySS [v2.3.4] (Jackman et al. 2017). Length distributions were calculated using samtools [v1.9] (Danecek et al. 2021) and graphed using R [v4.0.2] (R Core Team 2020) with the tidyverse package (Wickham et al. 2019). Minimap2 [v2.17-r941] and the pafCoordsDotPlotly R script (https://github.com/tpoorten/dotPlotly) were used to create dot-plots. For the Cypriniformes data-set, only chromosome level assemblies were included in the dot-plots. The *Danio rerio* (zebrafish) and *Triplophysa tibetana* genomes were also excluded from the dot-plots since few of the alignments passed the default quality filter in pafCoordsDotPlotly. BUSCO with the actinopterygii_odb10 database was used to find the BUSCO scores for each genome. The annotated genes from this new assembly were also compared to all annotated Cypriniformes using OrthoFinder [v2.5.4] (Emms and Kelly 2019).

**Table 1:** List of Cypriniformes genomes used in comparative analyses. The “L” column is an abbreviation of the assembly level: (S)caffold and (C)hromosome

| Organism Scientific Name | Assembly Name | Assembly Accession | L | Contig N50 | Size | Submission Date | Gene Count |
| --- | --- | --- | --- | --- | --- | --- | --- |
| *Anabarrilius grahami* | BGI_Agra_1.0 | GCA_003731715.1 | S | 36.06 Kb | 991.89 Mb | 2018-11-15 | 23,906 |
| *Carassius auratus* | ASM336829v1 | GCF_003368295.1 | C | 821.15 Kb | 1820.62 Mb | 2018-08-09 | 83,650 |
| *Cyprinus carpio* | ASM1834038v1 | GCF_018340385.1 | C | 1.56 Mb | 1680.12 Mb | 2021-05-12 | 59,559 |
| *Danionella translucida* | ASM722483v1 | GCA_007224835.1 | S | 133.13 Kb | 735.30 Mb | 2019-07-22 | 35,803 |
| *Danio rerio* | GRCz11 | GCF_000002035.6 | C | 1.42 Mb | 1373.45 Mb | 2017-05-09 | 40,031 |
| *Onychostoma macrolepis* | ASM1243209v1 | GCA_012432095.1 | C | 10.81 Mb | 886.57 Mb | 2020-04-17 | 24,754 |
| *Pimephales promelas* | EPA_FHM_2.0 | GCA_016745375.1 | S | 295.77 Kb | 1066.41 Mb | 2021-01-24 | 26,150 |
| *Puntigrus tetrazona* | ASM1883169v1 | GCF_018831695.1 | C | 1.42 Mb | 730.80 Mb | 2021-06-10 | 40,303 |
| *Sinocyclocheilus anshuiensis* | SAMN03320099<br>.WGS_v1.1 | GCF_001515605.1 | S | 17.27 Kb | 1632.70 Mb | 2015-12-14 | 52,005 |
| *Sinocyclocheilus grahami* | SAMN03320097<br>.WGS_v1.1 | GCF_001515645.1 | S | 29.35 Kb | 1750.27 Mb | 2015-12-16 | 55,200 |
| *Sinocyclocheilus rhinoceros* | SAMN03320098<br>_v1.1 | GCF_001515625.1 | S | 18.76 Kb | 1655.77 Mb | 2015-12-14 | 53,875 |
| *Triplophysa tibetana* | ASM836982v1 | GCA_008369825.1 | C | 2.83 Mb | 652.93 Mb | 2019-09-12 | 24,398 |

### ddRAD-seq sample collection and library prep

Fin clips were taken from the founders of the WorldFish Rohu Genetic Improvement Program, as described in Hamilton et al. (2019). A custom R script (https://github.com/IGBB/rohu-genome/) was used to minimize sampling putatively related founders (Hamilton et al. 2019). In total, fin clips from 64 male and 56 female rohu were sampled, sourced from the Halda (39), Jamuna (38), and Padma (43) rivers.

Genomic DNA was extracted from the samples using the Qiagen DNeasy Blood & Tissue Mini kit (Qiagen, Valencia, CA, USA). The concentration and purity of extracted genomic DNA samples were evaluated using a NanoDrop ND-1000 spectrophotometer (NanoDrop Technologies, Wilmington, DE, USA). The quality of genomic DNA was validated by electrophoresis on a 0.8% w/v agarose gel. The ddRAD-Seq libraries were made using the method described in Magbanua et al. (2022) with different restriction enzymes (NsiI and MspI) used to digest the genomic DNA and the adapters (Supplemental Table 2) in the ligation reaction. The libraries were submitted to Novogene (www.en.novogene.com) for a total of two lanes of PE150 Illumina HiSeq X-Ten (Illumina, San Diego, CA, USA) sequencing.

### SNP discovery and population analyses

Reads were mapped to the rohu genome using bwa [v0.7.17]. Single-nucleotide polymorphisms (SNPs) were called over two rounds using the Sentieon pipeline (Kendig et al. 2019) [Spack version sentieon-genomics/201808.01-opfuvzr] and following the DNAseq guidelines. Briefly, SNPs were predicted for the ddRAD-seq samples using the DNAseq pipeline for all samples, and these SNPs were used as known sites during base quality score recalibration (BQSR) in the second iteration of the DNAseq pipeline. The final SNP set was filtered via vcftools [Spack version 0.1.14-v5mvhea] (Danecek et al. 2011) to remove sites with insufficient representation (i.e., present in <90% of samples). Nucleotide diversity (π) and divergence (πxy, or dxy) were calculated in 10 kb windows using pixy v1.2.5.beta1 (Korunes & Samuk 2021) run via Miniconda3 [Spack version 4.3.30-qdauveb]. Population differentiation (Fst) was also calculated in pixy using 10 kb windows. Output from pixy was processed in R [4.1.1] and visualized using ggplot2 (Wickham 2016). Specific parameters and code can be found at https://github.com/IGBB/rohu-genome.

### Sex-Associated Fragments

To find regions of the rohu genome associated with sex, two-sample Monte Carlo tests comparing the high-quality read mappings for male and female samples were run for each fragment between the two digestion sites. The digestion site fragments for the rohu genome were found using egads (https://github.com/IGBB/egads). The high-quality read mappings for each sample were calculated by first filtering high-quality (mapq >= 30) mappings using samtools [v1.9] (Danecek et al. 2021), and then using the bedtools [v2.28.0] (Quinlan and Hall 2010) coverage to count the number of mappings to each fragment. Given the maximum selected size (613 bp) and the paired read size (300 bp), fragments with less than half of the sequence covered in a sample were removed from further analysis. Fragments with fewer than 50 samples (90% of the smallest sample group) surviving the filter were removed altogether. The fragment read mappings for each sample were normalized based on the total number of high-quality read mappings within a sample. Permutation tests were run on each fragment for 100,000 replicates, and the resulting p-values were adjusted using the Benjamini-Hochberg method. Fragments with an adjusted p-value less than 0.05 were considered associated with sex. The commands and code used can be found at https://github.com/IGBB/rohu-genome/y-link.

## Results & Discussion

### Genome Size Estimation

The C-value of rohu was previously reported as 1.99 pg (~1.95Gb) based on Feulgen densitometry (Patel et al. 2009) and 1.5Gb using k-mer estimation (Das et al. 2020); however, our flow cytometry results based on five individuals and our k-mer-based genome size estimation suggest that the rohu genome size is 50-65% the size previously reported by Patel et al. (2009) and Das et al. (2020), respectively. Our flow cytometry results indicate a C-value of 0.99 pg (~0.97Gb) with a standard deviation of only 0.02 across all measurements (Table 2). Moreover, our k-mer-based estimate using GenomeScope is 0.97Gb, the same value determined by our flow cytometry analysis. Lastly, our final genome assembly size for rohu is 0.95 Gb. Notably, the Feulgen densitometry estimate reported in Patel et al. (2009) for a second fish, *Labeo catla* (synonymous with *Catla catla*), was also approximately twice that later reported (Sahoo et al. 2020), perhaps suggesting stochastic differences, including cryptic variation in ploidy and/or differences in measurement techniques (Greilhuber 2005).

**Table 2:** Flow cytometry results for 5 rohu blood samples, all except Rohu-1 measured twice.

| Specimen Name | Number of Sample nuclei | Average sample fluorescence | Number of standard nuclei | Average standard fluorescence | Estimated Genome size<sup>2</sup> | HAPLOID |
| --- | --- | --- | --- | --- | --- | --- |
| Rohu-1 Rep 1 | 16020 | 27350 | 2065 | 69247 | 2.05 | 1.02 |
| Rohu-2 Rep 1 | 13082 | 25929 | 6570 | 66671 | 2.02 | 1.01 |
| Rohu-2 Rep 2 | 15402 | 25665 | 4354 | 67489 | 1.97 | 0.99 |
| Rohu-3 Rep 1 | 15124 | 25798 | 4442 | 68195 | 1.96 | 0.98 |
| Rohu-3 Rep 2 | 14923 | 25763 | 4823 | 68837 | 1.94 | 0.97 |
| Rohu-4 Rep 1 | 13320 | 26346 | 5913 | 69665 | 1.96 | 0.98 |
| Rohu-4 Rep 2 | 5624 | 26612 | 4097 | 68876 | 2.01 | 1.00 |
| Rohu-5 Rep 1 | 6771 | 25761 | 3080 | 68825 | 1.94 | 0.97 |
| Rohu-5 Rep 2 | 15926 | 26369 | 3352 | 68832 | 1.99 | 0.99 |
| Standard only<sup>1</sup> | 3 | 25258 | 3311 | 64331 | NA |  |
|  |  |  |  | **Average** | 1.98 | 0.99 |
|  |  |  |  | **Standard Deviation** | 0.04 | 0.02 |

<sup>1</sup> Trout erythrocyte nuclei: Genome size = 5.19pg.
<sup>2</sup> Genome estimate calculated as (average sample fluorescence/ average standard fluorescence \* standard genome size) in picograms DIPLOID (i.e., 2C).

### Genome Assembly & Annotation

Genome assembly was started with (a) a total of 130.5 Gb of Nanopore data, derived from 44.7 million reads, (b) 261 Gb of Illumina short reads (870 million 150 bp paired-end reads), and (c) 382 million 150 bp paired reads (114 Gb) from a Hi-C library. The initial *de novo* assembly was generated using the Nanopore data and polished with the short insert Illumina data, resulting in 4,999 contigs with an N50 of 1.28 Mb. After the Bionano and Hi-C data were incorporated, the total number of sequences dropped to 2,899 and the N50 increased to 29.9 Mb. These sequences were ordered and oriented by RagTag using the *Onychostoma macrolepis* reference to produce a final assembly with 25 chromosome-length scaffolds (deemed Chr01 through Chr25 - Supplemental Table 3) and 2,844 unplaced scaffolds, which ranged in size from 1,479 bp to 7.18 Mb. The chromosome scaffolds were composed of one to eight sequences each, with all but three composed of three or fewer sequences. The final assembled genome size was 945.5 Mbp, representing 97.9% of the estimated genome size (see Table 3 for assembly statistics at each step).

**Table 3:**
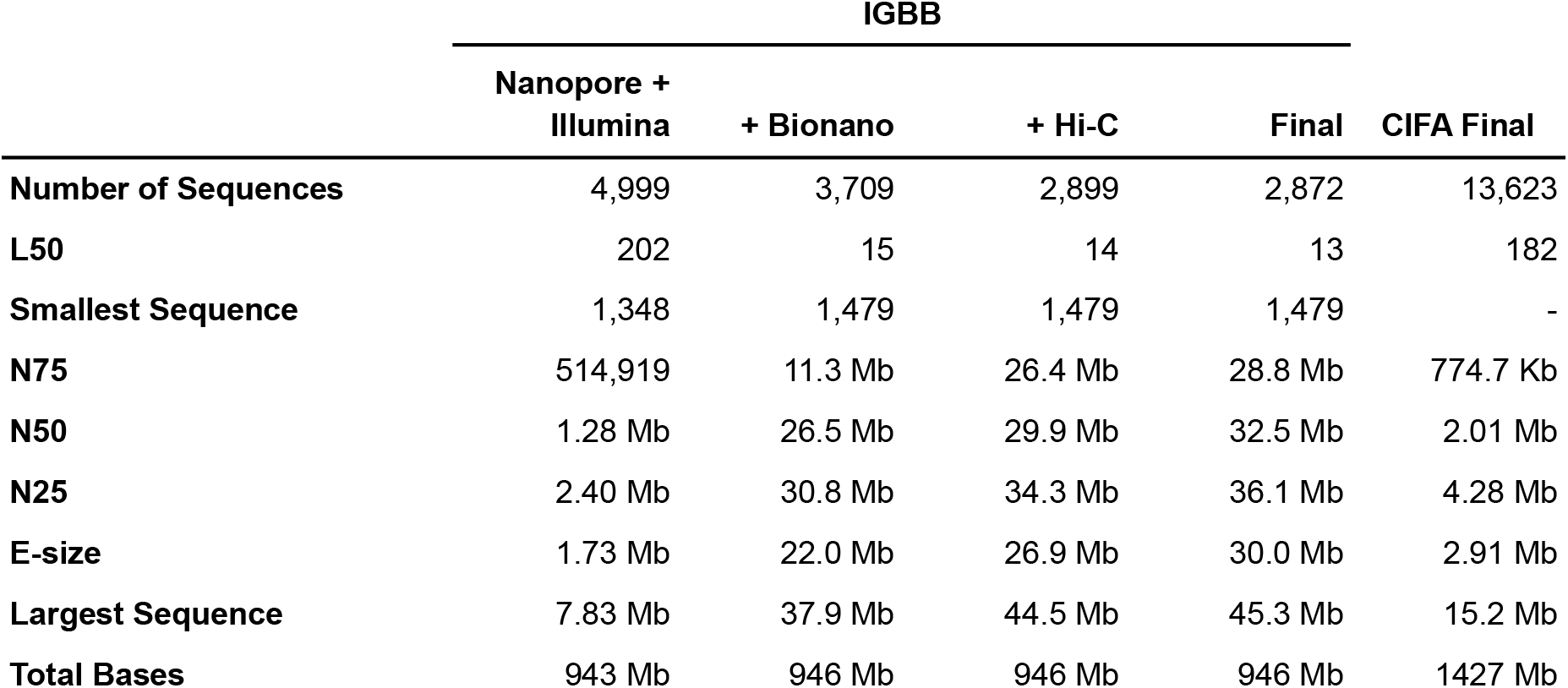
Assembly statistics for each stage of the IGBB rohu assembly and the CIFA rohu assembly.

RepeatMasker masked 41.25% of the genome. Due to the *ab initio* nature of the repeat database created by RepeatModeler2, most (92.5%) of the repeats found were unclassified.

The annotation pipeline identified 51,079 primary transcripts, of which 31,274 survived the AED, gap, and overlapping filter criteria. BUSCO analysis shows the genome includes complete copies of 98.1% of the 3640 orthologs in the actinopterygii_odb10 database with 37 (1%) duplicated. The filtered transcriptome contains 84.5% of the total orthologs complete with 74 (2%) duplicated. An overview of the BUSCO analyses can be found in Table 4.

**Table 4:** BUSCO analysis for the genome and transcriptome, before and after AED filtering.

| Type | Genome | Unfiltered<br>Transcriptome | Filtered<br>Transcriptome |
| --- | --- | --- | --- |
| Complete BUSCOs (C) | 3571 | 3139 | 3078 |
| Complete and single-copy BUSCOs (S) | 3534 | 3064 | 3001 |
| Complete and duplicated BUSCOs (D) | 37 | 75 | 74 |
| Fragmented BUSCOs (F) | 23 | 192 | 170 |
| Missing BUSCOs (M) | 46 | 309 | 392 |
| Total BUSCO groups searched | 3640 | 3640 | 3640 |

### Comparative genomics

Our assembly (IGBB) was compared with the published and publicly available rohu assembly (CIFA), and annotated Cypriniformes assemblies from NCBI that were scaffold level or higher. Both the scaffold N50 and maximum length of the IGBB assembly are 30 Mb longer than the CIFA assembly (Table 3). The length distributions (Supplemental Figure 1) show a similar separation, with overall greater contiguity in the IGBB genome. Interestingly, when the two rohu assemblies were pairwise aligned and plotted (Figure 1), the CIFA assembly shows a few large gaps, specifically in Chr09 and Chr19, despite being larger in size. Due to the twofold size difference between the assemblies and the fragmentation of the CIFA assembly, the inverse comparison (i.e., IGBB aligned to CIFA) was not informative. Dot-plot alignments of the chromosome level Cypriniformes assemblies (Supplemental Figure 2) generally exhibited similar chromosome structures, with some duplications and/or rearrangements. The assemblies for *Danio rerio* and *Triplophysa tibetana* were removed from the dot plot grid since very few of the alignments passed the graphing threshold. Comparing the BUSCO results for the rohu assemblies, the IGBB assembly had fewer duplicate, fragmented, and missing BUSCOs than the CIFA assembly. Furthermore, the IGBB assembly had the most single-copy BUSCOs of any Cypriniformes (Figure 2), even surpassing the model fish *Danio rerio.* Notably, *Carassius auratus* and *Cyprinus carpio* are both allotetraploid fishes (Xu et al. 2019; Braasch 2020) and therefore exhibit a good deal of duplication in the dot-plots and BUSCO results. Lastly, the annotations for the Cypriniformes were compared using OrthoFinder. Of the 31,274 genes annotated, 29,904 (95.6%) were placed into 18,740 orthogroups, which comprise 63.5% of the total orthogroups found. Supplemental Table 4 contains the summary statistics for all species used in the OrthoFinder analysis.

**Figure 1:**
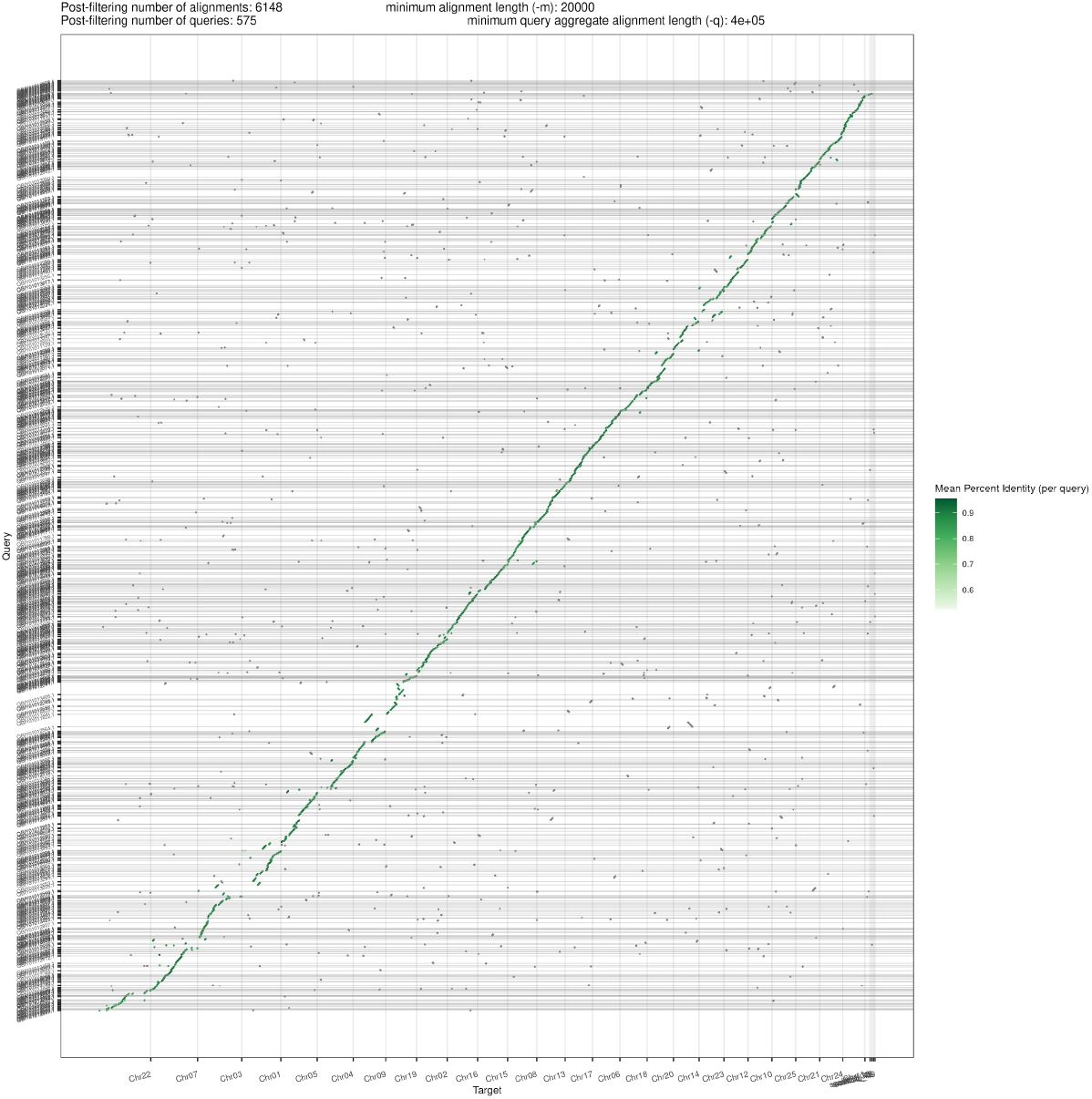
Dotplot between CIFA (y-axis) and IGBB (x-axis) rohu genomes, plotted using pafCoordsDotPlotly (https://github.com/tpoorten/dotPlotly)

**Figure 2:**
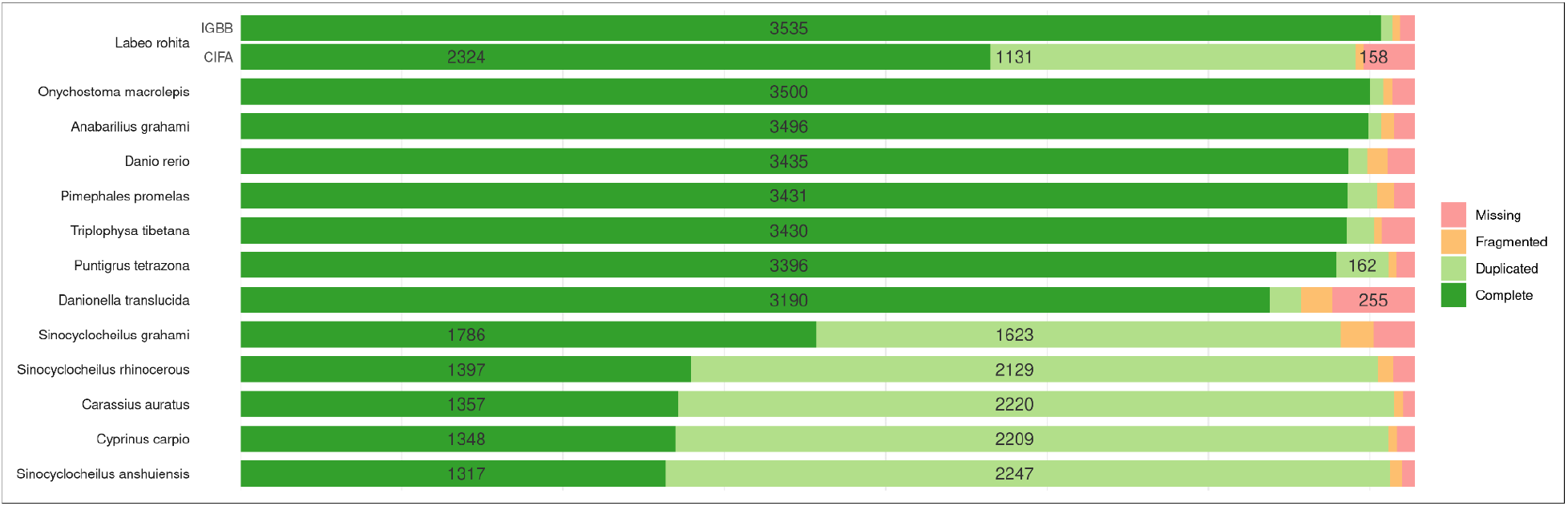
BUSCO results for both rohu genomes (IGBB and CIFA) and the other included Cypriniformes genomes. The results for the two groups are sorted by complete single-copy BUSCOs.

### SNP discovery and population similarities among rohu fisheries

Aquaculture is an agricultural growth industry, producing 46% of the fish consumed worldwide. Over 50 million tonnes of finfish are raised in aquaculture each year, with the vast majority of aquaculture occurring in Asia (FAO 2020). Farm-raised rohu comprises 3.7% of the finfishes produced annually and represents the 11th most commonly farmed finfish (FAO 2020). Consumer preferences have been surveyed, identifying traits (e.g., length and weight) to prioritize in improvement programs (Mehar et al. 2022) along with disease resistance, some of which may be multigenic and complex. Genetically improved rohu seed is increasingly available to farmers (Hamilton 2019; SZA 2021 Jun 10) (Das Mahapatra et al. 2007; Rasal et al. 2017; Hamilton et al. 2019; Hamilton et al. 2022); however, there is interest in further improving the characteristics of farmed rohu. Here we used ddRAD-sequencing in conjunction with the reference genome to provide insight into diversity and divergence among rohu in the Halda, Jamuna, and Padma river systems.

Patterns of divergence between the river systems (Figure 3A; Supplemental Table 5; Supplemental Figure 3) suggest that the geographically proximal Padma and Jamuna river systems (the Jamuna flows into the Padma) exhibited far less differentiation than either does to the hydrologically isolated and geographically distant Halda river system. While this pattern is similar to what was observed with silicoDArT markers (Hamilton et al. 2019), the greater number of nuclear sites surveyed here (i.e., 1.4 million) suggests that the differentiation between fish inhabiting these river systems is somewhat greater than previously reported using < 2000 SNP sites (Supplemental Table 5). These results (i.e., low differentiation between Padma and Jamuna and greater differentiation than previously reported) are congruent with an analysis of population structure (k=2) that reveals similar profiles for Padma- and Jamuna-based fish and a more divergent profile for fish from the Halda river system (Figure 3B). Interestingly, population structure analysis reaches optimization for these fish at k=1 (Figure 3C), possibly indicating greater than expected admixture between Halda fish and those from the other two, geographically distant rivers (Figure 3A).

**Figure 3:**
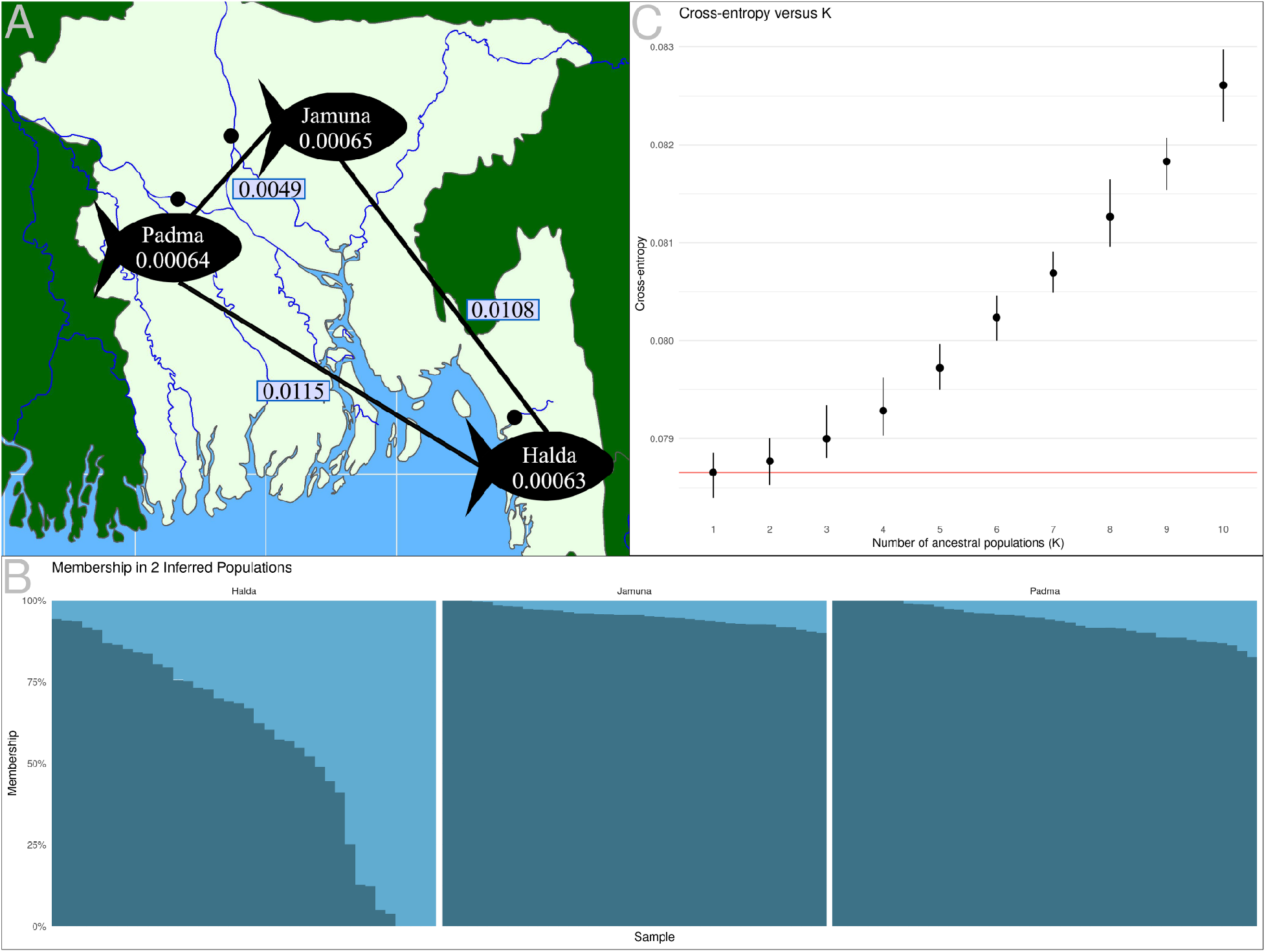
(A) Map of the river locations (dot), diversity within each population (number within fish), and divergence between each population (number between fish); (B) LEA predicted population structure (k=2) separated by river of origin. The vertical columns show the proportion (Q) assigned to each population for each individual; (C) Cross-entropy summary for the LEA analysis, using K=1 to 10 with 10 repetitions each.

Diversity among fish within each river was remarkably similar, ranging from 0.00063 in Halda to 0.00065 in Jamuna (Supplemental Table 6; Supplemental Figure 4). Notably, these estimates were nearly identical to the estimates of between-population divergence (πxy Supplemental Table 7; Supplemental Figure 5), which was 0.00064 for Padma-Halda and 0.00065 for both Padma-Jamuna and Jamuna-Halda, possibly indicating that these river populations are still representative of their shared ancestry. Diversity within populations (π) and divergence between (πxy) populations were distributed relatively evenly across the chromosomes; however, in both cases, chromosomes 3, 4, and 22 were the only chromosomes that exhibited greater than average π and πxy, possibly indicating differences in selection and/or permeability on those chromosomes. Interestingly, while Fst for chromosomes 3 and 4 were not considerably different from many of the other chromosomes, chromosome 22 exhibited the greatest relative population divergence (Fst; Supplemental Table 5), perhaps indicative of biologically relevant phenomena.

### Sex-Associated Fragments

The genetics underlying sex determination in fish can be complicated and variable even within species (Devlin and Nagahama 2002; Volff et al. 2007; Parnell and Streelman 2013; Heule et al. 2014; Nguyen et al. 2021); however, controlling the sex ratio is essential to optimizing farming of finfish (Martínez et al. 2014). Rohu breeding, for example, requires specific environmental conditions (i.e., monsoon (Natarajan and Jhingran 1963; Qasim and Qayyum 1962)), which is currently circumvented using hormonal induction (Bhattacharya 1999).

Despite its importance to aquaculture, the mechanisms governing sex-determination in rohu are currently unknown. Karyotypic analyses suggest that if rohu has sex chromosomes, they are likely homomorphic (Bhatnagar et al. 2014), similar to many other fish (Heule et al. 2014), and are indistinguishable from the remaining chromosomes. We screened the rohu genome for regions linked to sex by evaluating read mapping in each ddRAD region from female versus male fish. Between 9.8 and 23.4 million (M) reads were uniquely mapped per sample to the 473,345 genomic regions occurring between the two restriction sites, as predicted by egads. Approximately 42% of these regions (200,543) had at least 50 samples with >50% of the region covered and were retained for two-sample Monte Carlo testing. Monte Carlo testing highlighted 25 fragments from three chromosomes/scaffolds (Chr25, scaffold_1958, and scaffold_971) as significantly (BH adj. p-value <= 0.05) different between females and males with respect to read coverage (Figure 4, Supplemental Table 8). Interestingly, the seven significant fragments on Chr25 are (1) contiguous, (2) cover approximately 30 kb (26,052,217 - 26,083,955), and (3) have no female samples mapping, suggesting that this may be a male-specific region of chromosome 25. The five fragments on scaffold_1958 show a similar pattern, albeit with a shorter total length (6.1 kb) and with female reads present, although significantly diminished, for a single region of the scaffold (~100 bp). Conversely, both male and female samples map to the 13 significant regions of scaffold_971 with reasonable coverage; however, the female samples generally had around double the mapping rate relative to the male samples, suggesting that this region may be represented by a greater copy number in females versus males. Together, these results suggest that rohu has a male-heterogametic (XX/XY) system of sex determination. Furthermore, since the sex chromosomes are indistinguisable by karyotype (Bhatnagar et al. 2014) and the uniquely male regions comprise only a small region of the chromosome, rohu may only have a Y-specific region (or “young Y”), similar to *Oryias latipes* (medaka) (Kondo et al. 2006); however, sequences similar to the medaka homolog for male-determination (i.e., dmY (Matsuda et al. 2002; Hornung et al. 2007)) were not found in the rohu assembly, indicating that further study is needed.

**Figure 4:**
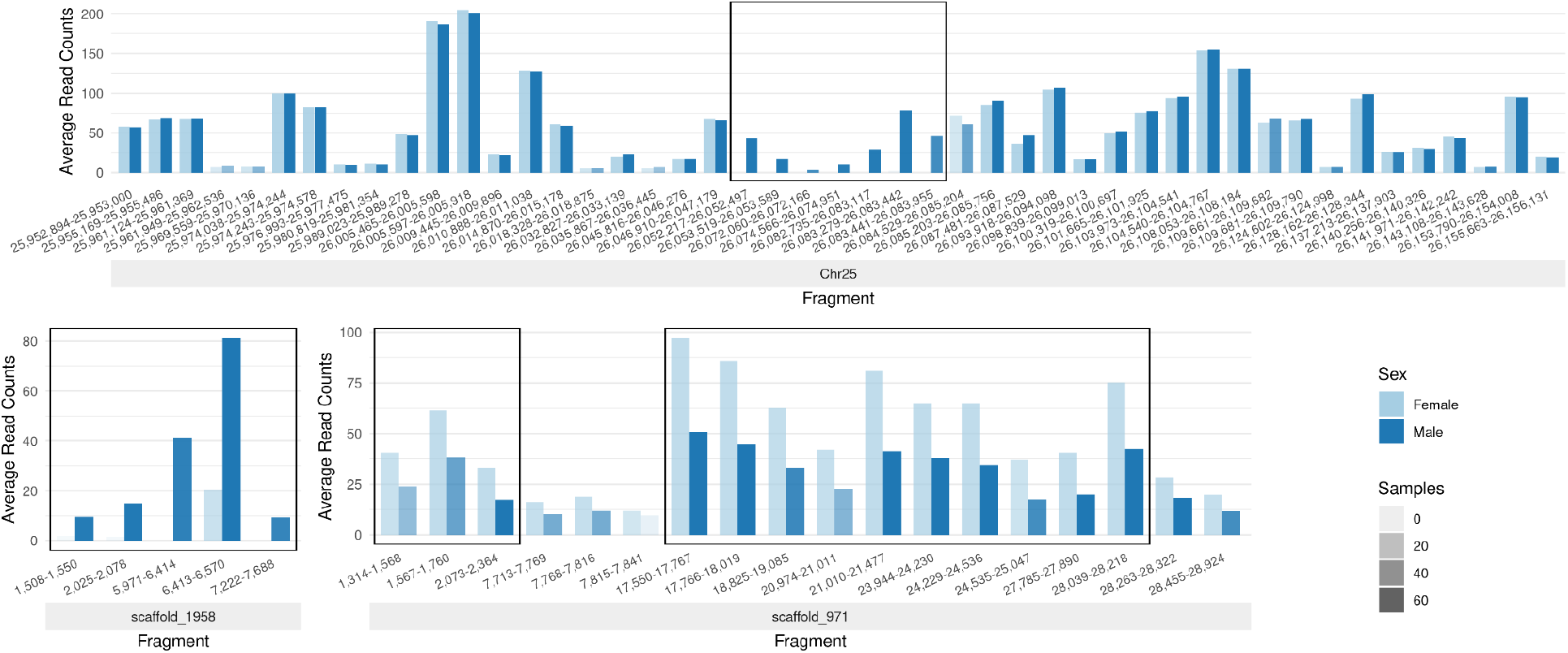
Regions of statistically significant differences between male and female rohu read counts. Each pair shows the average read counts for male and female samples for a ddRAD fragment. Opacity of the bars vary with the number of samples present. The fragments are ordered but not spaced according to position. Contiguous statistically significant fragments are outlined.

## Conclusion

Despite its importance to aquaculture, rohu has only recently been studied using modern molecular techniques. Our flow cytometry, kmer analysis, and next-generation sequencing assembly of the rohu genome indicate a genome size for *Labeo rohita* of 0.97Gb, a size 50-65% smaller than previously reported. Our IGBB reference-quality genome for rohu both improves contiguity and removes the excessive redundancy of the previously existing CIFA draft genome sequence. The IGBB reference genome is a valuable resource for breeding programs and evolutionary biologists as demonstrated in our initial ddRAD-seq experiments. We find that, while fish from the connected rivers (Jamuna and Padma) are more similar in relative divergence, there remains a question of whether these populations are recently diverged from the Halda river system or if there remains some gene flow between all three, despite the hydrological and geographical isolation of the Halda. We also report candidate regions for sex determination in rohu that may underlie a male-heterogametic (XX/XY) system. While greater sampling will be required to understand the genetics underlying rohu sex determination and the population dynamics of these river systems, the present information provides a foundation for breeders to facilitate aquacultural improvement of rohu.

## Data Availability Statement

The data used for the rohu genome and annotation are available at NCBI under the BioProject PRJNA650519. The assembled genome sequence and annotations are available at GenBank under accessions JACTAM000000000. The raw data is available at the SRA (Sequence Read Archive) under accessions SRR12580210 - SRR12580221. The ddRAD-seq data used for SNP discovery, population analyses, and sex-associated fragment analysis are available under the BioProject PRJNA841581 and the SRA accessions SRR19358298 – SRR19358417.

## Acknowledgements

The authors thank the Iowa State University Flow Cytometry Facility and ResearchIT unit for technical and computational support, respectively, WorldFish Carp Genetic Improvement Program staff based in Jashore, Bangladesh for managing and sampling fish, and Mahirah Mahmuddin for sample collation and management.

